# Modelling of the Cardiopulmonary Responses to Maximal Aerobic Exercise in Patients with Cystic Fibrosis

**DOI:** 10.1101/155713

**Authors:** Craig A. Williams, Kyle C. A. Wedgwood, Hossein Mohammadi, Owen W. Tomlinson, Krasimira Tsaneva-Atanasova

## Abstract

Cystic fibrosis (CF) is a debilitating chronic condition, which requires complex and expensive disease management. Exercise has now been recognised as a critical factor in improving health and quality of life in patients with CF. Hence, cardiopulmonary exercise testing (CPET) is used to determine aerobic fitness of young patients as part of the clinical management of CF. However, at present there is a lack of conclusive evidence for one limiting system of aerobic fitness for CF patients at an individual patient level.

Here, we perform detailed data analysis that allows us to identify important systems-level factors that affect aerobic fitness. We use patients’ data and principal component analysis to confirm the dependence of CPET performance on variables associated with ventilation and metabolic rates of oxygen consumption. We find that the time at which participants cross the anaerobic threshold (AT) is well correlated with their overall performance. Furthermore, we propose a predictive modelling framework that captures the relationship between ventilatory dynamics, lung capacity and function and performance in CPET within a group of children and adolescents with CF. Specifically, we show that using Gaussian processes (GP) we can predict AT at the individual patient level with reasonable accuracy given the small sample size of the available group of patients. We conclude by presenting future perspectives for improving and extending the proposed framework.

Our modelling and analysis have the potential to pave the way to designing personalised exercise programmes that are tailored to specific individual needs relative to patient’s treatment therapies.

## INTRODUCTION

Cystic fibrosis (CF) is the most common life shortening genetic disease in the Caucasian population, affecting nearly 11,000 individuals in the United Kingdom (UK) (1). The pathology of the disease, for which there is no cure, manifests itself throughout the respiratory, digestive and reproductive systems of the human body. Currently, as there is no cure for CF, the management of the disease is a key factor in the quality of care and health related quality of life factors. Part of the management of this disease requires cardiopulmonary exercise testing (CPET) to determine aerobic fitness, as represented by the maximal oxygen consumption (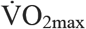). This parameter provides a clinically useful prognostic evaluation of a patient’s functional capabilities. Even in mild to moderate severity of CF, patients are known to demonstrate impairments in cardiac and respiratory functions leading to exercise intolerance.

Enhanced aerobic fitness has been shown to improve quality of life in young patients with CF (6-18 years), with benefits including lower risk of hospitalisation, increased exercise tolerance, reduced residual volume, increased endurance of the respiratory muscles, enhanced sputum expectoration and decreased rate of decline in pulmonary function (2–6). It has also been shown from CPET that individuals with CF possessing a higher 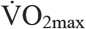 are shown to have a reduced mortality risk. Nixon *et al.* (1992) reported that individuals with a 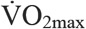 greater than 82% of their predicted value had an 83% 8-year survival rate, compared to just 28% 8-year survival rate for patients with a 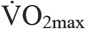 less than 58% of their predicted value (7). Furthermore, patients with a higher 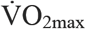 also have additional benefits in terms of improved fluid balance, retention of serum electrolytes through increased plasma volume, and potential for impact on sweat gland function thus reducing thermal strain and dehydration (8). These systemic changes are advantageous responses to exercise training, and result in an enhanced quality of life, increased physical function and increased life expectancy (9, 10).

Effective management of the disease has become even more critical in recent years due to an aging CF population group, with the median predicted survival of children born with CF in the UK now being 45 years (1). As a consequence of an aging patient group and high medical care costs (11), maintaining or enhancing fitness is crucial. Exercise has being widely acknowledged as a key management strategy for CF, supported by some mechanistic data on the systemic effects of exercise at the cellular level in vivo in young patients with CF (5-9). However, an integrated systems level understanding of the limitations of aerobic fitness for CF patients is lacking. Measurement techniques that do exist to quantify within-organ, real-time perfusion and intracellular oxygenation are invasive and unethical for use with paediatric patients, and current animal model research provides limited direct relevance to paediatric pathology. In clinical practice, there is significant interaction between cardiac and pulmonary function and the behaviour of the systemic vasculature during exercise training. This can result in the functional improvement in one part of the combined system, but detrimental effects on others (9). Clinicians therefore inevitably have to adopt very imprecise guidelines related to exercise prescription (12).

The use of modelling and simulation tools in clinical medicine is currently the subject of intense research interest both in the UK and internationally (13-17), and the adoption of a systems biomedicine approach to build and validate novel multi-scale, organ-level, integrated, re-usable and re-deployable models represents a paradigm shift in biomedical modelling and simulation. There are numerous organ level models in existence (18-21), however, to date there have been limited attempts to either integrate these or to apply them to real clinical applications. There is ongoing basic science and clinical trial work providing data on the micro (22) and macrovascular (23) changes associated with exercise. These data, although important, have yet to be integrated quantitatively with other data streams. In particular, there has been almost no previous work on the use of predictive modelling and simulation technologies for developing treatment strategies for CF patients.

Here, we present detailed data analysis of responses to progressive exercise in patients with CF, with a view of determining predictors of performance. We find that the time at which participants cross the anaerobic threshold (AT), as measured by means of gas exchange threshold (GET) is well correlated with overall performance. To gain further insight, we then develop a surrogate (statistical) model that allows us to evaluate how CF impairs exercise tolerance relative to increasing ventilatory and metabolic demands. Our modelling and analysis was based on data collected during cycling exercise form a CPET at different work rates (from resting to voluntary exhaustion) in young patients with CF. The outputs produced are discussed in this paper, with analyses focussing on pulmonary parameters.

## METHODS

Before turning our attention to modelling, we perform an exploratory analysis of the available data in order to identify predictors of performance. This study was carried out in accordance with the recommendations of the European Respiratory Society, written consent and assent was obtained from parent(s)/guardian(s) and participants, respectively. All participants’ parents gave written informed consent in accordance with the Declaration of Helsinki. The protocol was approved by the South West NHS Research Ethics Committee. Data were collected from 15 children and adolescents with CF, who performed a valid (24) and reliable (25) combined ramp incremental and supramaximal (Smax) CPET to determine 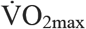 and the GET. This protocol was performed on an electronically braked cycle ergometer, and required patients to perform an initial exhaustive ramp incremental test at a pre-determined rate between 10–25 W⋅min^−1^, in order to elicit exhaustion in approximately ten minutes (26). After a 3-min warm-up at 10-20 W, participants completed this incremental test to the point of volitional exhaustion, maintaining a cadence of 70–80 rpm throughout. Exhaustion was defined as a 10 rpm drop in cadence for five consecutive seconds, despite strong verbal encouragement. Active (5-min cycling at 20 W) and then passive seated recovery (10 min) then preceded the Smax bout. Smax verification consisted of a 3-min warm-up (10-20 W), followed by a ‘‘step’’ transition to a constant work rate corresponding to 110% peak power output (27). Upon volitional exhaustion (defined previously), a 5-min active recovery (slow cycling at 20 W) concluded the combined CPET session.

## MODEL SIMULATIONS

Simulations are widely used in various fields of science and engineering because conducting physical experiments is too costly, or highly time-consuming, or even impossible in some cases (28). In the case of CPET in CF patients, there are also ethical considerations, since the test adds to the treatment burden many children and adolescents with CF already face.

Often, a primary goal of using model simulations is to perform quantitative studies such as uncertainty quantification or sensitivity analysis. Such studies are crucially important in biomedicine, since there exists significant variation both between and within patient groups. Through understanding and quantification of the uncertainty within the mathematical models, outcomes of patient-specific interventions can be better predicted. However, such investigations require a large number of runs that makes it impractical if each run takes more than a few seconds. To cope with this difficulty, one can use *emulators,* also known as *surrogates,* or *metamodels* or *response surfaces* (29). These provide a fast approximation of the input/output relation governed by the underlying simulator. The most important classes of surrogate models have been described elsewhere (30-32).

The surrogate model employed in this study is based on Gaussian processes (GP), which have become increasingly popular over the last decade (29). GPs have been used in a wide range of applications from wireless communication, to obtain position estimates for a mobile user (33); metallurgy, to model the development of microstructure (34); and in biology, to describe gene regulatory processes and cell growth (35-37).

The data analysis was performed using Python (Anaconda Software Distribution. Version 2-2.4.0. Continuum Analytics, 2016. URL https://continuum.io) and MATLAB and Statistics Toolbox Release 2016b, The MathWorks, Inc., Natick, Massachusetts, United States. The GP model simulator was implemented in R (R Core Team (2013). R: A language and environment for statistical computing. R Foundation for Statistical Computing, Vienna, Austria. URL http://www.R-project.org/.)

## RESULTS

### Data Analysis

To facilitate understanding, we first plot in **Figure 1(a)**, raw data displaying the performance of the participants. The work rate for each participant is increased at a rate that is either, a) dependent on their performance in previous tests, or b) when a prior test is unavailable, at a rate that is predicted to elicit exhaustion in approximately ten minutes (26). This is done in order to keep the expected duration of the test comparable to other participants. Note that this means that the total energy expended by a given participant is not based on the duration of the test alone. In **Figure 1(b)**, we show how participant age affects overall test performance. We observe a correlation between the two: the worst performing participants tend to be the youngest, but this effect is insignificant at older ages. The colour coordination used in this figure (red-worst performance → blue-average performance →green-best performance) will be used throughout the remainder of this section, where performance is quantified by the total energy transferred during the test.

**Figure 1:**
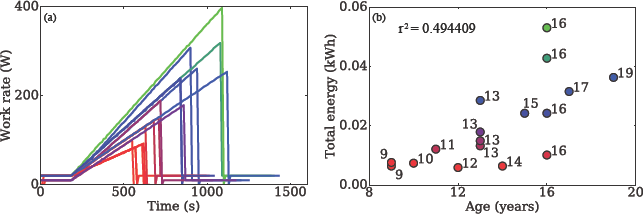
(a) The work rate for each participant is increased at a rate dependent on their past test performance. (b) Participant age is correlated with test performance for young participants, but not for older ones.

In **Figure 2**, we plot the ratio of 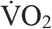 over total ventilation 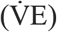 with respect to time. The markers on each of the time traces indicate the time of volitional exhaustion for that participant. There are two features that stand out from this figure. Firstly, participants who perform better have higher 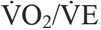 ratios, suggesting that their oxygen uptake is more efficient than their poorer performing counterparts. Secondly, in the recovery phase of the test (5 minutes following volitional exhaustion) better performing participants exhibit a sharp decrease in 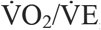, which is not observed in the poor performance group. Again, this suggests a more efficient utilisation amongst the former group and that exhalation of CO_2_ is perhaps more significant to total breathing following the test.

**Figure 2:**
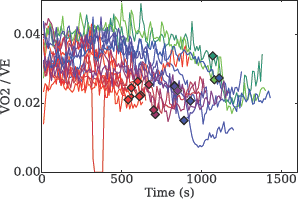
Ratio of oxygen utilisation and total breathing throughout the test. Markers indicate the volitional exhaustion times for each participant.

We next examine the effect that breathing patterns have on participant performance. Two classical prognostic measures used for patients with cystic fibrosis are the forced vital capacity (FVC), and the forced expiratory volume in one second (FEV_1_). These measures have been shown to be well correlated with mortality and overall fitness of CF patient groups (38–40). In **Figure 3**, we demonstrate how these metrics are correlated with performance in the CPET test. In **Figure 3(a)**, we observe good correlation between FVC and the maximum tidal volume (TV) of breathing achieved throughout the test. This is unsurprising since participants are likely to be trying to maximise their breathing depth close to their exhaustion point. However, notice that, although the group with low FVC performed poorly, this measure was unable to separate other participants. **Figure 3(b)** reiterates this result and also highlights the high correlation between FEV_1_ and FVC.

**Figure 3:**
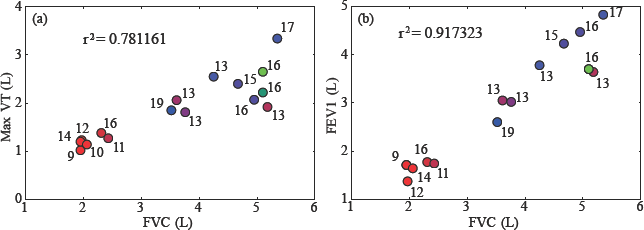
(a) Correlation of FEVi with the maximal tidal volume achieved throughout the test. (b) Correlation between FEV_1_ and FVC is high. Note that, although FVC and FEV_1_ are good predictors of poor test performance, they are unable to distinguish better performing participants.

In order to better classify the performance of the participants, we must instead look for other factors. In **Figure 4**, we present the total breathing rate against the oxygen consumption throughout the test. In **Figure 4(a)**, we find a strong relationship between test performance and respiratory pattern. Note that the curvature of the graphs suggests that an exponential fit, rather than a linear one, is most appropriate for these data. In order to test this, we take logarithms of the data and perform a linear regression, ignoring the first 180s of the test since participants are here in the warm up phase (work rate is not increasing) and the final 60s of the data prior to volitional exhaustion, since participants pass their respiratory compensation point, inducing hyperventilation and erratic breathing. The results of the fit are shown in **Figure 4(b)** and we can see more clearly the association of performance on breathing pattern.

**Figure 4:**
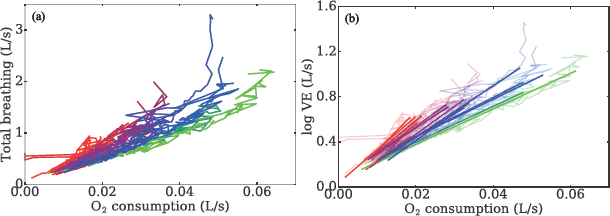
(a) Total ventilation plotted against oxygen utilisation. We observe that breathing pattern is strongly correlated with test performance. (b) Exponential curves are fitted through the raw data, further highlighting this dependence.

From the fitted curves, we can further explore the dependence of breathing patterns on performance. Firstly, in **Figure 5(a)**, we plot the slope of the fitted curve against the total energy transfer. We find that the slope of the curve of 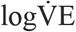 against 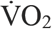 alone does not capture all of the variation in energy, which is highlighted by the low R-squared value (0.68). Instead, we plot in **Figure 5(b)** the oxygen consumption at a fixed rate of breathing against the total energy. Here, we find a good characterisation of the overall performance, with a much higher R-squared value (0.86), confirming that those who utilise oxygen more efficiently perform better.

**Figure5:**
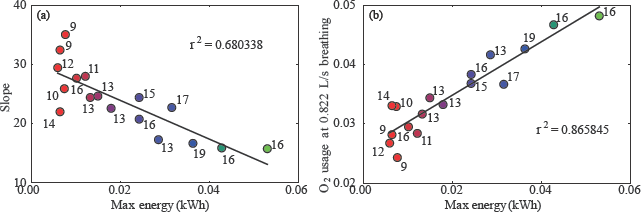
Slope of the fitted curves (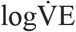 against 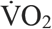) from Fig. 4(b) plotted against the total energy transfer during the test. We find a relatively poor characterisation of the variance between performances. (b) By instead plotting the oxygen consumption at a fixed rate of breathing, we better capture differences in performance.

Next, we examine the specific patterns of breathing exhibited by the participants, in particular, focussing on breathing depth and frequency. Initial characterisations of these patterns seem to provide little information, as indicated in **Figure 6(a)**. However, when we now also include dependence of oxygen consumption, we find a near perfect classification of participants into the lowest performing groups, the best performing groups and the middle group. These data are displayed in **Figure 6(b)**. Note that in this figure, the trajectories appear to be evolving on a planar manifold, suggesting significant co-dependence between these three variables.

**Figure6:**
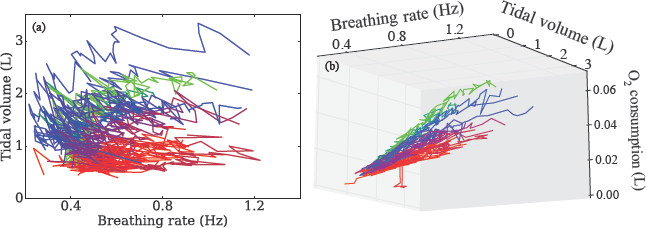
(a) Breathing patterns subdivided into the breathing rate and tidal volume. These data appear uninformative for predicting test performance. (b) With the additional inclusion of the oxygen consumption at a fixed rate of breathing, we find that these variables now almost perfectly capture variation in participant performance.

Given that there appears to be co-dependence between the variables used in **Figure 6(b)**, a sensible next step is to use principal component analysis (PCA) to account for these dependencies. By projecting the data onto their principal components, we show in **Figure 7** how well these capture the variation in participant performance. Given that there are only three independent variables in our analysis, it is natural to use spherical polar coordinates to show how these quantify performance. The first of these components, θ, captures over 90% of the variation in performance (**Figure 7a**), as does the normal component in the direction of breathing frequency (**Figure 7b**). These results further indicate the importance of breathing frequency, together with co-variation of oxygen consumption and tidal volume as predictors of test performance.

**Figure 7:**
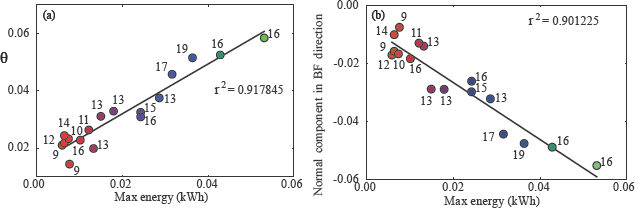
First of the principal components obtained via PCA accounts for over 90% of the variation in test performance. (b) Similar levels of variance are accounted for by taking only the normal component of the first principal component in the breathing frequency direction.

### Gas Exchange Threshold

Under steady state levels of exercise, the metabolic rate of production of CO_2_ is assumed to be proportional to the utilisation rate of O**2** via the cellular respiratory quotient, since (after the initial rest-work transition) adenosine triphosphate (ATP) is replenished primarily via aerobic metabolism pathways. As the work rate increases, this pathway becomes unable to supply sufficient ATP to satisfy the required amount of energy and anaerobic pathways have to contribute to overcome the shortfall. In doing so, they increase the levels of waste products, such as lactate and also increase the overall production rate in CO**2**. The point at which this occurs is known as the anaerobic threshold (AT) or sometimes referred to as the lactate threshold. However, this is difficult to directly measure *in vivo* during exercise, and as such the GET is utilised as a non-invasive surrogate of the AT (41).

The time at which participants cross the anaerobic threshold is well correlated with overall performance, as shown in **Figure 8(a)**, due to the fact that the anaerobic pathways are less efficient at producing ATP and because build-up of lactate contributes significantly to fatigue. One of the major contributing factors in defining AT is 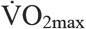, since this is indicative of the limit of the rate of oxidative phosphorylation. It thus comes as little surprise that 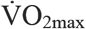 is the best single predictor of CPET performance, as shown in **Figure 8(b)**.

**Figure 8:**
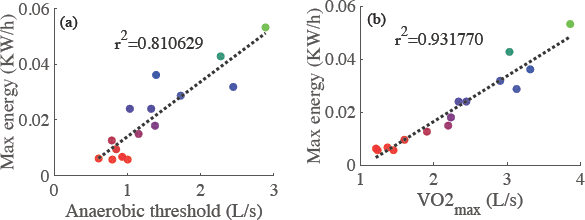
(a) Quantifying the relationship between the anaerobic threshold and overall test performance. Anaerobic thresholds were calculated using an automated procedure based on previous methods (41) (b) V O_2max_ is the best single predictor of overall test performance.

### Gaussian Processes-based Modelling

A Gaussian Process (GP) defines a probability distribution over functions where the true function is considered as a particular sample path. In our modelling, we attempt to describe the influence of breathing patterns and 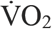 on the AT, since this is shown to correlate well with overall test performance (see **Figure 8**). In mathematical terms, we treat AT as our scalar output variable, with input variables comprising: baseline breathing rate and tidal volumes, and O_2_ consumption rates at a fixed ventilation rate, FVC, FEV_1_, and the rate of changes in breathing rate and tidal at exercise onset, using the slopes calculating in **Figure 4(b)**. Thus, we have an output variable dependent on seven input variables, which we conveniently store in a vector **x** ∈ *D* ⊂ ℝ^*d*^, with *d=7.* We now assume that there exists a ‘true’ function *f: D →* ℝ, such that AT = *f*(**x**).

A GP is fully specified by its mean, μ, and covariance *K,* which are both functions of the input variables: µ = µ(**x**), *K* = *K*(**x,x’**). Specifically, if *Y* is a GP, then we write:

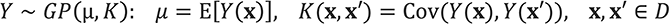

The above can be regarded as prior distribution over function spaces. This can be seen more clearly in **Figure 9(a)**. In this subfigure, which shows a generic example of a GP, the bold red line is the ‘true’ function *f* Note that the true function is unknown – our aim is to construct a model that approximates it. The thin grey lines are sample paths of a GP with a zero mean prior and a constant prior covariance function. For exposition purposes, the example plot is restricted to the case *d*=1, but the approach is unchanged for *d* > 1. In principle, μ; could be any function, for practical purposes, polynomial regressions are common choices. The choice of covariance function reflects our prior belief about the structure of *f* (such as the level of smoothness) and therefore has a crucial influence on our modelling. A typical choice for the covariance function is the squared-exponential kernel, given by:

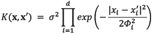

Here, σ controls variability of *Y* along the y-axis while *Φ*_*i*_ > 0, *i* = 1, … , *d*, scale the distance measure for each input dimension. A more thorough exposition of common choices for the covariance function may be found in (42).

Thus far, we have defined the prior distribution for the GP. It is clear from Figure 9(a) that, in general, prior distributions are unlikely to provide a good approximation to the true function *f*. We can incorporate data (or evaluation of *f* at specific points), with our prior distribution to give a posterior distribution, following a Bayesian framework. The resulting posterior distribution of the GP, conditioned on the data, will be much closer to the true function.

Let **y = {*f*(x^1^), …,*f*(x^n^)**} be a set of function evaluations at *n* locations **X = {x^1^, …,x^n^**}. Here, function evaluations correspond to the AT location for a set of patients during the CPET. Predicting with GP is obtained by conditioning *Y* on sample points Ω = (**X, y**}. For any (new) **z** ∈ *D*, the posterior distribution of *y*(**z**)|Ω has a normal distribution with the following mean, *m,* and variance, *s^2^*

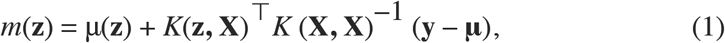

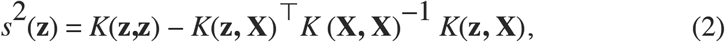

Where ^T^ denotes the transpose operator, ^-1^ is the inverse operator and **μ** = μ**(X)** is the vector of the mean function at **X**. In addition, K(**z,X**) and *K*(**X,X**) are the covariance vector between *Y*(**X**) and *Y*(**Z**) and the covariance matrix between the observations. Figure 9(b) shows an example of incorporating sample points to update the prior distribution shown in Figure 9(a). In this generic example, the function *f* is evaluated at five distinct values of x, and the mean and variance of the GP are updated using (1)-(2).

**Figure 9:**
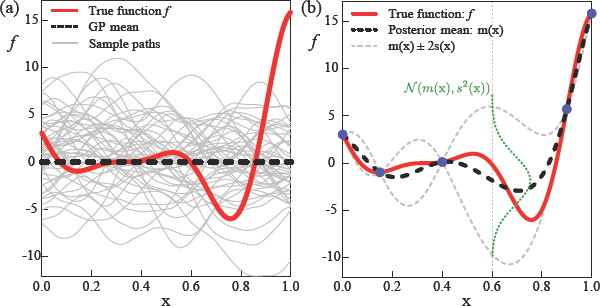
GP is a probability distribution on function spaces. The thick solid red line is the true function *f* and the thick dashed black line is the GP mean. (a) Thin grey lines show sample paths of a prior GP whose mean is zero. (b) Blue bullets indicate five data points sampled from *f*. The GP mean and covariance are updated using these sample points. The thin grey dashed lines show *m(x)* ± 2s(x).

At the evaluated points, indicated in blue (colour online), the true value of ***f*** is known and so the variance of the GP at these points vanishes and µ(**x**) = *f*(**X**). In between these points, the variance increases, dependent on the distance from an evaluated point. The mean of the GP, shown by the thick black dashed line now approximates the true function much more closely (recall that the prior mean function in Figure 9(a) was zero everywhere), and matches exactly at the evaluated points. The approximation can be further improved by incorporating more data (function evaluations), particularly around those input values for which the variance is high. Thus, as more data becomes available, the model is iteratively improved.

### Justification of Variables

In our simulator, we have used the AT as our output (dependent) variable. Another choice for this could be the performance in the CPET or 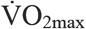, since these are the primary biomarkers for gauging aerobic fitness. However, the use of the GET has been shown to have high agreement with the lactate threshold (another surrogate for the AT), and related to disease severity in CF (43). Furthermore, as reported in **Figure 8(a),** the AT location for a given patient correlates well with their overall performance in this test. Furthermore, by constructing a predictive model to approximate the AT values for a patient, we can hope to further extend this to identify contributions of aerobic and anaerobic pathways in supplying ATP to meet the demand imposed during the exercise test.

The initial exploration of results highlighted that both ventilation parameters and metabolic rates of O_2_ consumption were the primary factors influencing test performance. It is clear that V O_2_ should play a significant role in determining the AT location, since it is a proxy for oxidative phosphorylation which is the main pathway for ATP synthesis in steady state exercise. As a measure of oxygen uptake efficiency in our model, we use the oxygen consumption rate at a fixed total ventilation rate (that being 0.822 L/s) as an input (independent) variable for each patient.

There are a number of ventilatory input variables incorporated in our simulator. Given their potential importance as clinical biomarkers, highlighting the limitations of lung capacity and function, we include FVC and FEV_1_ as input (independent) variables. During the aerobic exercise test, participants spend three minutes cycling at a minimal work rate, over which we quantify their baseline breathing frequency and baseline tidal volume by taking the means of these variables over this period. To capture the dynamics response associated with the exercise, the rates of change of breathing frequency and tidal volume are calculated, based on the fits obtained in **Figure 4(b)**. The rates of change of these ventilatory variables indicate how participants respond to changes in workload and were shown in **Figure 6(b)** to discriminate between participant performances. Moreover, differences in rates of change of breathing frequency and tidal volume have previously been shown to be significantly different between control groups and CF groups (44), suggesting that these are potentially key biomarkers for assessing aerobic fitness in patients with CF.

### GP Emulator Performance

The GP emulator was constructed using the data presented above, with AT calculated using previously described methods (41). These data were used to train the emulator. For the prior distributions, a first order polynomial regression was used for the mean, whilst a squared-exponential kernel with σ and Φ provided via maximum likelihood estimation was used as the prior covariance. Since in this pilot study, we have a small number of participants, we use leave-one-out cross-validation to assess the accuracy of our emulator, that is, for each patient, we use the emulator trained against the remainder of the sample points to approximate the AT value for that patient, given their input variables. The results of this are presented in **Figure 10(a)**, in which we also plot the 95% confidence intervals, with relative percentage errors demonstrated in **Figure 10(b)**.

**Figure 10:**
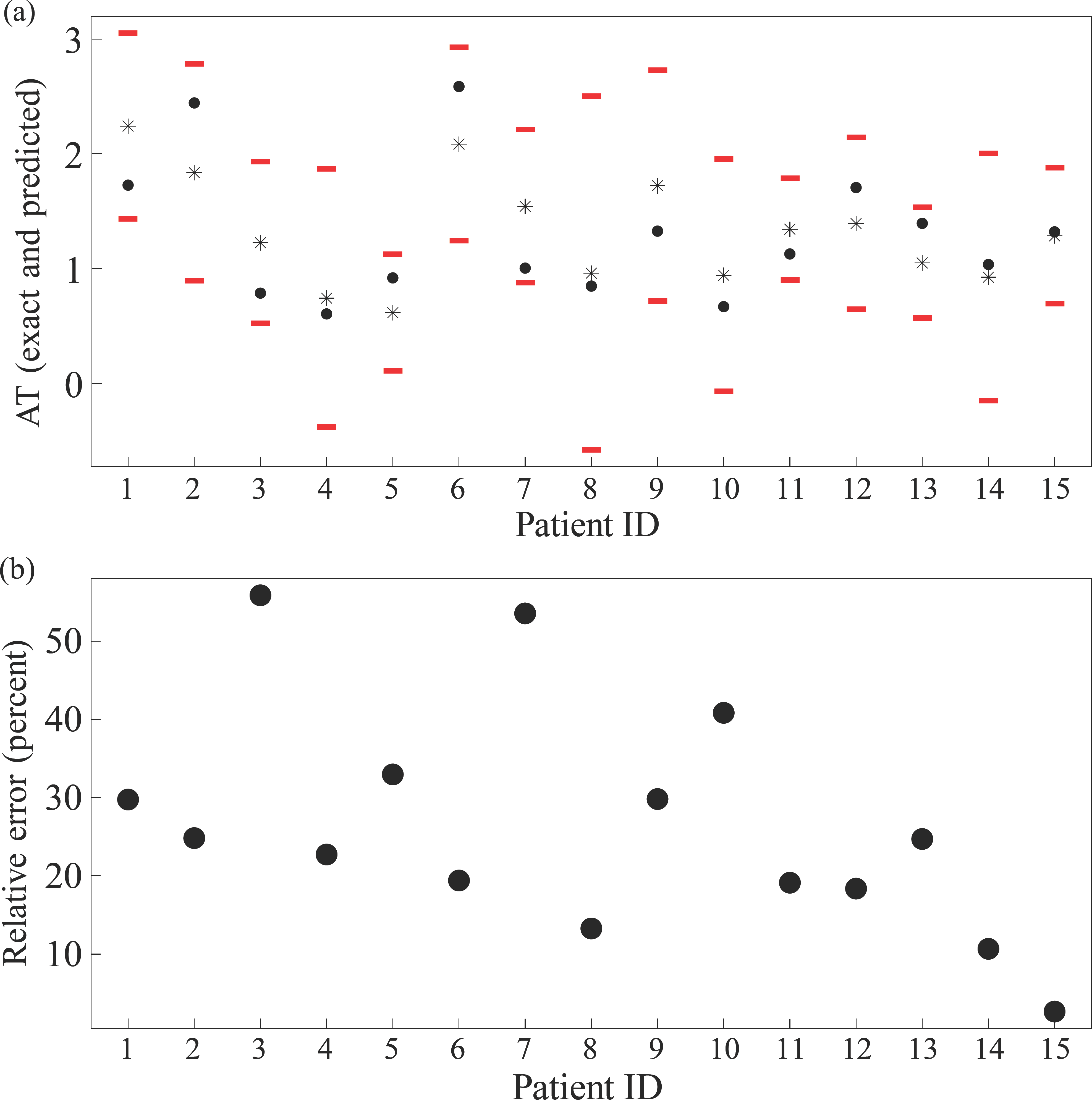
(a) Predictions (asterisks) vs. exact values (bullets). Red bars show the 95% confidence intervals, *m*_i_(x_i_) ± 1.96 *s_i_*(x_i_) around the predicted value, where *m*_i_ and s_i_ are GP prediction mean and standard deviation based on all but the i’th data point. (b) Relative prediction error expressed as a percentage of the true value *f*(x_i_).

The emulator has reasonable accuracy, in spite of the small sample size. In general, for high accuracy in GP emulator, the number of sample points (corresponding to the number of patients in our case) should be at least ten times larger than the number of input variables (45). Clearly, there is a need to acquire further data points to improve the predictive capabilities of the emulator. For each patient, the 95% confidence interval around the predicted point contains the true value. Moreover, the relative errors for many patients is small (considering the low sample size), though we note that some patients, (3 and 7), the error is high. This highlights a need to extend this study to include more data to refine estimates around these points, particularly to deal with the high variability of lung function parameters in this patient group (46).

## DISCUSSION

Our primary modelling aim is to eventually use the model to evaluate how CF impairs exercise tolerance relative to increasing ventilatory and metabolic demands. Our predictive model could also be used to evaluate therapies and their effect on exercise performance. Ultimately, we hope that this will form a series of steps to design better exercise treatment that is tailored to specific individual needs relative to patient’s treatment therapies, a treatment modality that is affordable, and personalised (47).

The data analysis and modelling results have highlighted the dependence of CPET performance on variables associated with ventilation and metabolic rates of O_2_ consumption. Whilst these observations are, in themselves, not novel, we believe that this is the first attempt to mathematically model the relationship between ventilatory dynamics, V O_2_ and performance in CPET within a group of children and adolescents with CF. Whilst it is clear that there is much work to be done in this area, we hope that this will serve as a starting point for improved modelling, not only in the arena of GP emulators, but also in the domain of mechanistic modelling, which we shall describe briefly.

### Perspectives for GP improvements

At present, the GP model is conditioned on specific data points for each patient. An improvement to the GP could be made by instead conditioning with respect to distributions. Given that repeated tests are often performed for the same individual, so that multiple sample points are provided for each participant, we can consider a fit to a probability distribution capturing the variability in the identified variables. This approach has advantages compared to standard GP models, such as avoiding problems associated with overfitting and regularisation (which is important for the inverting ill-conditioned covariance matrices that often arise during the application of (1)-(2), (48).

### Perspectives for mechanistic modelling

In order to better understand and characterise the difference between performances, it would be extremely useful to construct and simulate a mechanistic mathematical model, based upon on ordinary differential equation (ODE) framework, describing the relationship between the cardiopulmonary system and the metabolic dynamics of skeletal muscle. By describing the relationships between different organ-level systems, the model would be able to identify the patient-specific rate-limiting factors defining aerobic fitness. Moreover, analysis of the model could be used to suggest treatment strategies to improve these factors and thus predict how patients will improve under such regimes.

At the individual organ level, there are a plethora of models describing individual dynamics of the level of the heart (49-52), lung (53-56) and systemic metabolic demand (57-63). There also exist a number of models describing such interactions between cardiopulmonary and metabolic systems (64-68) in a variety of settings, including heart failure and mechanical ventilation. A core feature in all of these models is the nonlinear interaction between the constituent model compartments that encompass the distinct tissues. An important consequence of this is that the model must be studied as whole, in an integrated fashion, to truly understand the body’s response to exercise.

With respect to the present question, there are a number of limitations of the existing modelling approaches. Most significantly, none have been designed with either an adolescent, or a CF patient group in mind, and the nuances of these patient groups will have to be factored into to any model development. In particular, these models have relatively simple, empirical models to describe changes in ventilation, which may not capture breathing dynamics of our patient group well. Moreover, to the best of our knowledge, no model considers the changes in ventilation separated into breathing frequency and depth that have been shown by us and others (44) to be critical to overall test performance.

In our analysis, we have demonstrated that the AT or GET location is a critical factor in determining overall patient aerobic fitness. Many of the mathematical exercise models describe only steady-state exercise, in which aerobic pathways meet most of the ATP demand (64-68). As such, these models are inadequate to capture the dynamics we describe here. Another common topic of study is the dynamic response at exercise onset, which again, does not meet the current need to describe the AT crossing point (69-72).

Of the mathematical models that describe the contribution of anaerobic pathways to ATP production, some assume that the shortfall in meeting ATP demand via oxidative phosphorylation is met entirely by anaerobic pathways (73), yet this is clearly not so, since ATP levels in skeletal muscle postexercise may be up to 30% lower than pre-exercise values (74). Mathematical models that factor in fatigue brought about by anaerobic metabolism are generally phenomenological in nature, and it is difficult to quantify these models against real patient data (73, 75, 76), and these models mostly fall outside the arena of ODE-based modelling and so dynamical properties are difficult to infer from them.

Developing a mathematical model to describe the integrated behaviour of all of the relevant organs, whilst remaining biophysically plausible, but without requiring excessive or invasive parameterisation is a difficult task. The proposed model should include descriptions of the cardiovascular system, the ventilatory system and simple models of metabolism at the tissue level. Specifically, dynamic variables should include alveolar, arterial, venous and tissue level partial pressures/concentrations of O_2_ and CO_2_, cardiac output, ventilation and metabolic rates oxygen utilisation and CO2 production. Partial alveolar gas pressures can be linked to data collected during the test, and the work rate can then be provided as inputs to the model. Note that these variables are similar to those included in previously defined models (64-68) and the aim is to extend these to describe the dynamics observed in patients with CF. The proposed model schematic is displayed in **Figure 11.**

**Figure 11:**
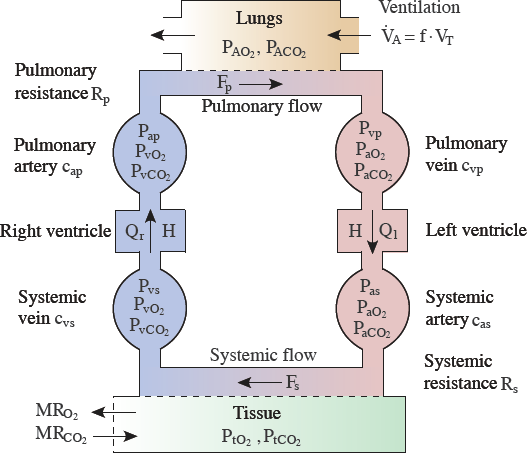
Schematic of the variables and processes in the proposed ODE-based mathematical model. Adapted from Timischl (1998) (66) and Batzel *et al.* (2005) (80).

Of critical importance to the overall model construction is the development of a simple, yet realistic model of cellular metabolism, to overcome the issues discussed earlier. The model should respect the different metabolic processes that occur in the muscle tissue, in particular: glycolysis, phosphocreatine breakdown and synthesis and oxidative phosphorylation, in a simplistic fashion that is amenable to being fit to CPET data. Whilst there are models that describe the biochemical reactions associated with these processes, and importantly, their stoichiometry (58, 61, 63, 67, 77), quantifying their associated rate constants *in vivo* is a near-impossible task, and so efforts must be made to develop a model that incorporates the relevant metabolic dynamics whilst being simple enough to be fit to data.

With knowledge of the integrated system, attempts can also be made to describe other important exercise-based processes, such as lactate buffering and recycling (as a fuel source) (61, 78, 79) and the overall muscle fatigue brought about by the combination of all of these factors. Only by systematically exploring the dependence of aerobic fitness of all of the factors described in this section can we begin to understand the system in an integrated fashion.

## LIMITATIONS

A limitation with the current study is the utilisation of a relatively small sample size, and this may be contributing towards aforementioned errors. Future studies should seek to utilise CPET collected annually in CF centres, to develop larger, multi-centre, samples whereby a uniform exercise protocol is utilised. Given that utilisation of CPET is now recommended and endorsed for regular use by international medical societies (81), and individual CF centres are reporting upon experiences of using CPET (82), large-scale utilisation of such data is a feasible target.

The findings presented may be derived from a smaller sample, and therefore the models presented are only preliminary results for this patient cohort; however, the study provides a unique examination into the aerobic and anaerobic signatures of individual patients with CF in response to progressive exercise.

Finally, whilst this study provides an insight into metabolic process during exercise, future research and models must account for additional variables predictive of function and mortality (e.g. genotype, body composition, pancreatic sufficiency, infection status, exacerbations (83, 84) and co-morbidities (CF-related diabetes (85), pulmonary arterial hypertension (86)) existent within CF, notably those that may affect exercise tolerance.

## CONCLUSIONS

Benefits of modelling include the ability to utilise existing data sets at a time when there are limited resources. There is also a call to reduce the burden on sick patients (EU directive). As models improve and the quality of fits to data are improved, the models can be used in a prognostic setting to predict potential improvements in aerobic fitness that may arise due to therapeutic intervention. Moreover, with proper mechanistic modelling of the primary organs affected in CF, there exists the potential to optimise treatment for this patient group by identifying the limiting factors of aerobic fitness.

## AUTHOR CONTRIBUTIONS

CAW conception, design of the work, data collection, analysis, interpretation, manuscript drafting, final approval of the version to be published, agreement to be accountable for all aspects of the work in ensuring questions related to the accuracy and integrity of any part of the work are appropriately investigated and resolved; KTA data interpretation, critical review of the paper, final approval of the version to be published, agreement to be accountable for all aspects of the work in ensuring questions related to the accuracy and integrity of any part of the work are appropriately investigated and resolve; OWT data collection, analysis, interpretation, final approval of the version to be published, agreement to be accountable for all aspects of the work in ensuring questions related to the accuracy and integrity of any part of the work are appropriately investigated and resolved; KW supervision of work organization and development, data analysis, interpretation, manuscript drafting, final approval of the version to be published, agreement to be accountable for all aspects of the work in ensuring questions related to the accuracy and integrity of any part of the work are appropriately investigated and resolved. HM construction of GP emulator, final approval of the version to be published, agreement to be accountable for all aspects of the work in ensuring questions related to the accuracy and integrity of any part of the work are appropriately investigate and resolved.

## ACKNOWLEDGMENTS

We are grateful to the participants for volunteering their time to this project, especially the young patients with CF. KW was generously supported by the Wellcome Trust Institutional Strategic Support Award (WT105618MA). KTA gratefully acknowledges the financial support of the EPSRC via grant EP/N014391/1.

## Supplementary Material

Supplementary Material should be uploaded separately on submission, if there are Supplementary Figures, please include the caption in the same file as the figure. Supplementary Material templates can be found in the Frontiers Word Templates file.

Please see the Supplementary Material section of the Author guidelines for details on the different file types accepted.

